# STED-FCS in subdiffraction limit volumes reveals altered diffusion in live cell applications

**DOI:** 10.1101/2025.10.23.684144

**Authors:** Bernd Zobiak, Maria Andres-Alonso, Yuanyuan Wang, Behnam Mohammadi, Christian Gorzelanny, Hermann Clemens Altmeppen, Shuting Yin, Antonio Virgilio Failla

**Affiliations:** UKE Microscopy Imaging Facility (UMIF), University Medical Center Hamburg-Eppendorf (UKE), Martinistraße 52, 20246 Hamburg, Germany; Leibniz Group "Dendritic Organelles and Synaptic Function", Center for Molecular Neurobiology Hamburg (ZMNH), University Medical Center Hamburg-Eppendorf, Hamburg, Germany; Department of Dermatology and Venereology, University Medical Center Hamburg-Eppendorf, Hamburg, Germany; Institute of Neuropathology; University Medical Center Hamburg-Eppendorf, Hamburg, Germany; Leibniz Institute for Neurobiology (LIN), 39118 Magdeburg, Germany

**Keywords:** Super resolution microscopy, STED-FCS, live cell imaging

## Abstract

Fluorescence correlation spectroscopy (FCS) is a widely established light microscopy technique for investigating physiological parameters such as diffusion states, particle numbers, and viscosity in biological samples. Combining FCS with stimulated emission depletion (STED-FCS) has enabled the investigation of molecular diffusion to sub diffraction-limited volumes. However, the full potential of STED-FCS for biomedical applications remains underexplored. Here, we present proof-of-principle studies with novel implementations of STED-FCS for investigating biological processes in living samples. Specifically, we demonstrate the impact of STED-FCS analyses in three distinct biomedical assays. Firstly, we prove that STED-FCS is capable of observing dynamic changes in autophagy-related protein microtubule-associated protein 1 light chain 3b’ (LC3b) along neuronal axons. Secondly, we show that STED-FCS can resolve alterations in the carbohydrate chain lengths of glycoproteins in melanoma cells. Finally, we demonstrate that STED-FCS can measure the reduced mobility of lipids within the plasma membrane of neuronal cells treated with the Alzheimer’s disease-associated, aggregation-prone toxic peptide amyloid-beta 1-42 (Aβ42). We believe that this study will inspire researchers to utilize STED-FCS to address critical questions in their bio-imaging studies, particularly regarding the super-resolution assessment of dynamic processes in living cells.

## Introduction

In biomedical research the elucidation of (patho-)physiological mechanisms in diseases, development requires a comprehensive understanding of underlying processes at nanoscopic levels. Currently, several light microscopy techniques are available for characterizing the physiochemical properties of biomolecules in model systems and living specimen, such as Förster resonance energy transfer (FRET) [1], fluorescence lifetime imaging (FLIM) [2], fluorescence recovery after photobleaching (FRAP) [3] or second harmonic generation (SHG) [4], and spectroscopic techniques like fluorescence correlation spectroscopy (FCS) [5], image correlation spectroscopy (ICS) [6], as well as Raman [7] or surface plasmon resonance (SPR) [8]. Notably, these techniques offer variable sensitivity, specificity and especially spatio-temporal resolution. In this contest, FCS turned out to be particularly advantageous for studying molecular diffusion with high temporal and spatial resolution. The basic principle relies on recording the time-dependent intensity fluctuations of freely diffusing fluorescent dyes within a defined volume. In the latter, diffusion can be influenced by multiple parameters associated with the local environment of the molecule, i.e., its own conformational state, molecular interactions, medium density and temperature [9]. Consequently, FCS allows researchers gain access to insights of the diffusion behavior of either single or multiple molecular species. This technique provides information to a wide range of parameters namely molecular concentration [10, 11], stoichiometry [12, 13], reaction, diffusion kinetic constants [14, 15] and viscosity [16]. The excellent biocompatibility of FCS for performing measurement non-invasively with living cells under physiological condition is a critical advantage when compared to the other widely used techniques, namely FRAP. Yet, the full potential of FCS is underutilized and typically only applied for studies performed either in solution or on isolated membranes such as supported lipid bilayers or giant unilamellar vesicles. Since biological systems are inherently complex in nature, comprising numerous regulatory and structural components, measurements within living cells are necessary for characterizing and studying molecules in their native environment.

The optical diffraction barrier, however, constitutes a major limitation in bioimaging experiments as it precludes observation in elipsoidal volumes of less than ∼200×200×700 nm³. In the case of confocal-FCS, such volumes might still enclose many molecules and are larger than, for instance, membrane nanodomains or multimolecular clusters [17]. Thus, some spatially confined changes in diffusion are challenging to be accessed, because they are likely to be averaged out. Combining FSC with STED microscopy has the potential to overcome the challenges mentioned above [18, 19]. Indeed, STED microscopy can generate an emission volume whose diameter is approximately 30-50 nm. This means that STED microscopy can probe fluorescence signals in focal regions at least 25 times smaller than those achieved with confocal microscopy. Since its first practical inception in 2009 [20], STED-FCS has been applied to a variety of biological studies. Some examples are the investigation of membrane reorganization of the bacterial pore-forming toxin listeriolysin O [21], nanodomain incorporation of GPI-anchored proteins in living cell membranes [17], and the role of the glycoprotein Env in the packaging of HIV-1 particles [22]. However, we believe that the potential of STED-FCS as a bioimaging tool to elucidate complex molecular processes in living systems is still not fully expressed. To this end, we will demonstrate three novel applications, exemplifying where STED-FCS brings deeper and new insight to the scientific observations: 1) Studying interactions of the autophagy-regulatory molecule LC3b in neuronal synaptic sites [23]; 2) measuring pathological alterations in the carbohydrate chain length of glycoproteins in a disease model; and 3) monitoring the influence of Aβ42 (the toxic, aggregating peptide implicated in Alzheimer’s disease (AD) pathogenesis) on neuronal cell membrane fluidity [24]. These novel applications offer valuable insights into disease mechanisms and further demonstrate the precise, quantitative capabilities of STED-FCS for characterizing dynamic molecular processes within biological systems.

Therefore, with these studies, the importance of STED-FCS as an imaging tool for live cell imaging applications at a nanoscopic resolution has been further demonstrated.

## Materials and methods

### Chemicals

Phenol red-free medium supplemented with 10% FCS was used for cell imaging. The other chemical reagents used in the experiments were SNAPCell 647-SiR (500 nM), DPPE-Abberior Star Red (0.25 mg/mL a.d.), WGA-Atto647N (1 mg/mL), rabbit-anti-goat IgG-Atto647N, LipofectamineTM 2000 (Cat.No. 11668027, Invitrogen), dimethylsulfoxide (DMSO, Cat.No. D8418-50ML, Sigma-Aldrich). A 2 mM stock solution was prepared by dissolving Aβ-FITC (synthetic ‘FITC-β-Ala-Amyloid β-Protein (1-42)’, Cat.No. 4033502, Bachem) in DMSO and aliquots were stored at -80 °C. 36 nm far-red fluorescent beads, Abberior solid mount with anti-fading reagent.

### Plasmids

LC3b-SNAP, SNAP-control, shRNA against EXT1 (shEXT1), shRNA-control (shCTL), GFP-Synapthophysin.

### Animals

Animals were housed in the ZMNH animal facility at the University Medical Center Hamburg-Eppendorf (UKE), Germany, under controlled environmental conditions. All animal experiments complied with the ethical guidelines for animal research outlined in the European Communities Council Directive (2010/63/EU) and were approved by both the Hamburg state ethics committee (Behörde für Gesundheit und Verbraucherschutz, Fachbereich Veterinärwesen) and the University Medical Center Hamburg-Eppendorf animal care committee.

### Cell culture

The cells were maintained in DMEM supplemented with fetal calf serum (FCS, 10%) and Penicillin-Streptomycin (50 units/mL) with an exception of mHippo cells, they were cultured without any antibiotics. Phenol red free medium supplemented with 10% FCS was used for cell imaging. Cells were maintained at 37 °C and 5 % CO2 throughout the experiments. Immortalized mouse embryonic hippocampal cells (mHippoE-14; #CLU198; CELLutions Biosystems Inc.) and MV3 cells were cultured in growth medium and routinely screened for mycoplasma contamination. Primary neurons were prepared from Wistar rat embryos of both sexes (E18). The neurons grow in neuronal growing medium containing BrainPhys (Stemcell Technologies, Cat no #05790) supplemented with 1X SM1 (Stemcell Technologies, Cat No #05711) and 0.5 mM glutamine (Gibco). They are plated onto a coverslip coated with poly-D-lysine at a density of 20.000–60.000 cells per well [25].

### Cell labeling

Neurons were seeded on 18 mm poly-L-lysine coated glass coverslips and co-transfected with GFP-Synaptophysin and LC3b-SNAP at 10-14 DIV according to the manufacturer’s instructions 2 days before the experiment. On the experiment day, cells were labelled with 0.24 µM SNAPCell 647-SiR substrate in growth medium for 15 min at 37 °C, washed twice with imaging medium and incubated another 15 min in imaging medium to reduce background labeling. Neurons were washed once with imaging medium and the coverslip was transferred to an imaging chamber (Warner Instruments, Cat No RC-49MFSH) filled with imaging medium for the measurement. MV3 cells were transfected with shEXT1 and control shCTL were performed as previously reported [26]. Cells were seeded in 8-well chamber slides (ibidi) and incubated at 37 °C and 5 % CO2 in a humidified incubator overnight until reaching 75% confluent. On the day of experiment, cells were labelled with 1 µg/mL of WGA-Atto647N in PBS at room temperature, washed three times with PBS and kept in PBS for the FCS measurements. mHippo cells were seeded in 35 mm glass bottom dishes (MatTek) one day before the experiment and treated for 2 h with Aβ-FITC (final concentration: 1 µM; controls were treated with DMSO), prior to the measurement at 37 °C. Cells were washed with cold PBS and labelled with 36 nM DPPE-Abberior Star Red in PBS on ice for 20 min. Cells were then washed three more times with cold PBS and kept in PBS for the measurement.

### Microscope setup

For all STED/confocal-FCS measurements, a commercial Abberior easy 3D STED (infinity line) system was used (Figure 4A). The wavelengths of pulsed lasers used for excitation were 488 nm and 640 nm, and for the pulsed depletion beam 775 nm (3 W). The beams were directed through a Nikon Ti microscope body equipped with perfect focus system and a 60x Na 1.4 P-Apo oil objective. A spatial light modulator formed the donut-shape of the depletion beam. The emitted light was detected through GFP (495/25 nm) and Far-Red (685/70 nm) filters placed in front of two independent avalanche photodiodes. The pinhole was set to 1 AU throughout all measurements. Abberior’s Imspector software was used to control the setup.

### STED-/FCS measurements

Cells were imaged either at room temperature (MV3 and mHippo cells) for slowing down internalization of the membrane-bound dyes, or at 37 °C (primary neurons). From an overview confocal scan of a single cell, 2-4 points were selected and fluorescence time traces recorded sequentially. In case of MV3 and mHippo cells, the axial position was set within the dorsal membrane of the cell. The axon of neurons is very thin, so that the focus was set at the maximum intensity level. For every point, the focus was stabilized using the system’s auto-focus. The laser beam was held at the selected position and the fluorescence signal continuously recorded for a certain amount of time, i.e., point scan FCS was performed for 60 s with a pixel dwell time set to 1 µs (xt mode). Minimal excitation power between 0.5-2 % was applied to reduce bleaching. For STED-FCS (Figure 4B), the depletion beam power was set to 20 % and the alignment checked on every measurement day to ensure equal detection volumes between the experiments. From the point spread function of sub-resolution 40 nm or 100 nm fluorescent beads, the beam waist in xy- and z-dimension (wXY and wZ, respectively) were calculated and used as a constant for the fit model (see section “data analysis” in the online Supplementary Methods).

### Data analysis

The further processing of spectroscopic raw data and quantifications are described in detail in the online Supplementary Methods. Briefly, the autocorrelation function (ACF) were computed from the fluorescence time traces and, depending on the characteristics of the diffusion and the dye, fitted according to different models (Figure 4B): 3D diffusion (equation 2 in online Supplementary Methods), 3D diffusion with two components (equation 3 in online Supplementary Methods) or 3D diffusion with triplet state extension (equation 4 in online Supplementary Methods), respectively. The diffusion time was determined and the diffusion coefficient calculated using calibrated beam diameters. Where applicable, the fraction of each component was determined. All values are summarized in Table S1 (online Supplementary Information).

## Results

### LC3 changes from a slow to a fast diffusing state after the entry from axons into synapses

Autophagy is a highly-conserved degradative pathway whose function relies in the formation of a double-membrane organelle called autophagosome that engulfs cargo. Amphisomes are intermediate organelles, formed during autophagy through the fusion between autophagosomes and endosomes [27]. In neurons, autophagosomes are known to form constitutively in distal axons [28, 29]. Recently, several reports showed that autophagy has acquired specialized functions in neurons [30, 31] and that synaptic activity is linked to autophagosome formation at synapses. However, whether and how autophagosomes are formed at synaptic sites following neuronal activity remains unknown. Light-chain 3b (LC3b) is a central protein and a marker of autophagosomes [32]. LC3b is a cytosolic protein that undergoes lipid conjugation to autophagosome membranes during the formation of the organelle and this is essential for autophagosome biogenesis. Despite the ubiquitous distribution of LC3b, it remains unclear whether the diffusion behavior of LC3b is affected by the molecular crowding at synapses, in particular presynapses, which would affect rapid autophagosome formation at these confined sites. One conclusive step to identify where these interactions occurs is to monitor LC3b’s diffusion coefficient using FCS in living hippocampal neurons. The experimental design consisted of seeding primary rat neurons on glass coverslips 48h before the experiment and, later, co-transfection with GFP-Synaptophysin (green, presynaptic marker) [33] and either LC3b-SNAP (red) fusion proteins or SNAP (red) as controls. In Figure 1A, a schematic of the experiment is depicted. Two regions of interest were selected, as presynaptic bouton site, and an adjacent region presynaptic boutons. The scheme suggests that, confocal-FCS acquisitions measures only mean diffusion for a large number of LC3b proteins, while detection capability of 2D STED-FCS is relatively fewer number of proteins. It is notable that the elements shown in Figure 1A are only for purpose of representation. In Figure 1B, the overall imaging workflow is illustrated. The sample overview is shown in confocal mode. Later, a region of interest was selected and simultaneously imaged with confocal and STED modes, respectively. The regions of interest selected for the FCS measurements were located inside the zone of presynaptic boutons (arrow, synaptic site) and the non-synaptic region is depicted with arrowheads. The four images in the upper row of Figure 1B were acquired of neurons expressing LC3b fused to SNAPCell647-SiR for detection while in the rest of the second row expression of SNAPCell647-SiR was used as a control. Autocorrelation curves (ACs) were plotted based on the obtained data from FCS experiments. In Figure 1C (top graph), examples of LC3b ACs are displayed. ACs of LC3b-SNAPCell647-SiR revealed two diffusing species that could be identified by a double-exponential fit (blue line) (equation 3, in online Supplementary Methods), as opposed to a single exponential fit (green line). The first diffusion component, called “fast “, is characterized by a diffusion time τ(d1) of <10 ms while for the second („very slow or static molecules “) component τ(d2) was >10 s. The double exponential fit allowed us to retrieve from each dataset the fraction of the fast and slow components. This information will be essential to fully analyze the results of this experiment. The “fast” diffusion time τ(d1) can be determined precisely by fitting the initial part (first 2s) of the AC with the mono exponential fit model (see an example in Figure 1C, middle panel). The mono exponential fit model is described in equation 2 (online Supplementary Methods). Experimental data suggested that diffusion coefficient of LC3b-SNAPCell647-SiR was measurably reduced in the analyzed region of interest while comparing measurements in confocal-FCS with STED-FCS. Thus, we further investigated the diffusion coefficient’s dependence on the size of the excitation spot by using three excitation spot diameters on the same location, the experiment conducted using “confocal mode” at circa 270 nm, “low STED” at circa 120 nm and “STED” at circa 75 nm (Figure 1C, lower graph). We observed that the reduction in diffusion coefficients at both synaptic and non-synaptic sites were proportional to the decrease in excitation spot size. This phenomenon is usually interpreted as anomalous diffusion [34], a process in which molecules are prevented to move due to external factors. As an important result, anomalous diffusion appears to be stronger in the synapses indicating a lower molecular mobility in these sites.

**Figure 1:**
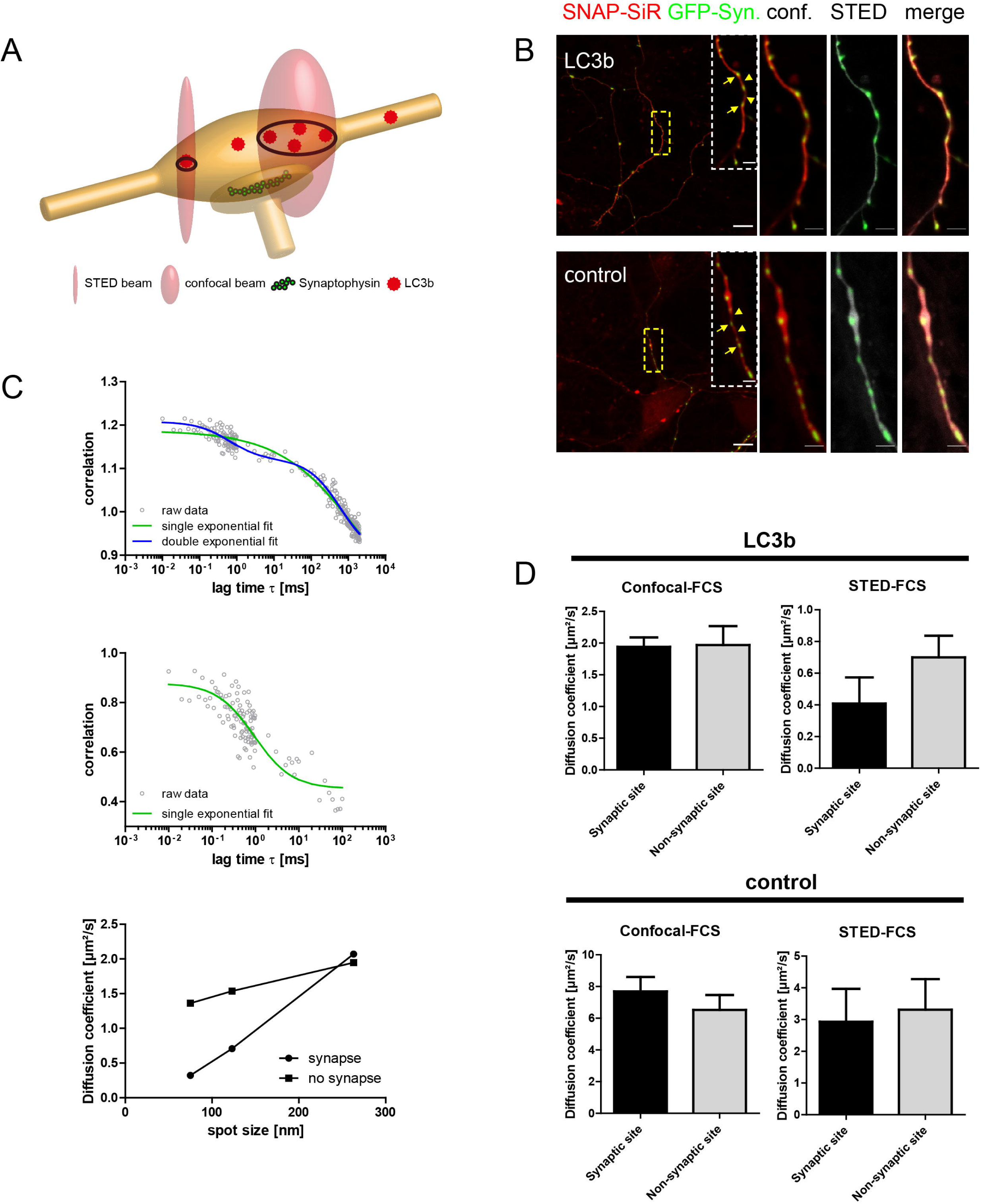
STED-FCS measurements of LC3b at synapses on rat hippocampal neurons. (A) Schematic view of the experiment: GFP-Synaptophysin (green) labels synaptic sites. FCS time traces from amphisomes (red) were detected in synaptic and non-synaptic site by means of confocal or 2D STED beam. Please note the draw elements are depicted not in scale to their real proportion. (B) From left to right: First frame, sample overview in confocal mode, i.e GFP-Synaptophysin (green), LC3b-SNAP (red). Second frame, region of interest acquired in confocal mode, i.e GFP-Synaptophysin (green), LC3b-SNAP (red). Third frame same region of interest: GFP-Synaptophysin (confocal, green), LC3b-SNAP (STED, gray). Fourth frame: overlay of frames 2 and 3. Scale bar: 10 µm, 2 µm (inset). (C) Top, autocorrelation curves of LC3b diffusion revealed two diffusing species identified by a double-exponential fit (blue line) according to equation 4, compared with a single exponential fit (green line). This 3D diffusion model revealed a fast (with diffusion times τ(d1) <10 ms), and a slower component (with diffusion times τ(d2) > 10 s). A precise determination of τ(d1) was achieved by applying a single exponential fit (green line) according to equation 2. The single component 3D diffusion model was applied lag times shorter of 10² ms of the autocorrelation curves. Bottom, diffusion coefficients of LC3-SNAP-SiR inside and outside synapses, were computed for different spot sizes of the excitation beam by tuning the power of the 775 nm STED beam. The slope of the curves revealed anomalous diffusion inside synapses. (D) Confocal-FCS and STED-FCS measurements were performed in the region of pre-synaptic boutons (Synaptic site) and outside of these regions (Non-synaptic site). The diffusion coefficient of the fast component [τ(d1)] was calculated for LC3b-SNAP-SiR (LC3b) and SNAP-SiR empty vector (Control). Top, while with confocal-FCS, no differences were detectable, in STED-FCS measurements the diffusion coefficient in synaptic sites was reduced about 50%. Also the fraction of the fast component decreased in synaptic sites by 30% (see Table S1). The differences were not apparent in the empty vector control, bottom figure. All values are harmonic mean ± SEM.

Furthermore, the data (Figure 1D) in the first graph of the second row show that confocal-FCS was unable to detect differences in fast diffusion coefficients between measurements performed in synaptic and non-synaptic sites. Importantly, we were able to detect a significant decrease of the fast diffusion coefficient within the synapses when we performed the recordings using STED-FCS, as depicted in Figure 1D. On the other hand, the results of confocal and STED acquisitions performed in the control sample failed to present any significant difference between the fast diffusion coefficients, irrespective of whether measured inside or outside synaptic sites (Figure 1D, bottom row). Together with the reduction of approximately 50% of the fast diffusion speed, STED-FCS measurements revealed a significant decrease of 30 % in the number of molecules that were diffusing within the synaptic sites (see Table S1). In summary, confocal-FCS measurements could not detect any significant variation in the average diffusion behavior of LC3b molecules between synaptic and non-synaptic sites. In contrast, the combination of STED-FCS was able to simultaneously detect a stronger anomalous diffusion in synaptic sites leading to a reduction in the diffusion coefficient as well as an overall reduction in the number of fast diffusing molecules.

### Variations in carbohydrate chain lengths of glycoproteins can be quantified in a model system

Further, STED-FCS is used for investigating carbohydrate chain length of glycoproteins. Previously, STED-FCS has been applied for studying the movement of the carbohydrate binding protein ‘wheat germ agglutinin’ (WGA) on the cell surface. The glycocalyx is a dense layer of variable carbohydrates that are present on the external cell-surface [35]. Heparan sulfate and hyaluronic acid are abundant on the cell surface and play a significant role as co-receptors for growth factors [36, 37]. Typically, chain lengths of carbohydrates are in the nanometer range under normal physiological conditions. It is widely reported that the chain lengths of glycocalyx components are reduced in tumor cells and certain diseases [26, 38–40]. Therefore, the ability to detect changes in lengths or in the assembly of the glycocalyx is important for clinical diagnostic applications. However, a reliable method to measure the variations of chain lengths is technically challenging [41]. The power of resolving nanoscopic volumes by STED-FCS microscopy provides a potential diagnostic tool in this regard. For this proof-of-principle demonstration, the human melanoma cell line MV3 was used as a model system. Based on our previous research, we modified the length of the heparan sulfate chains by shRNA-mediated (shRNA EXT1) knockdown of Exostosin-1 (EXT1), a gene involved in the biosynthesis of heparan sulfate [42], while control cells were only treated with shRNA (shCTL) [26]. In the above cases, cells were labeled with WGA-Atto647N, a lectin that binds to the carbohydrate chains of glycoproteins, including heparan sulfate. Figure 2A schematically describes the STED-FCS method applied for this experiment.

**Figure 2:**
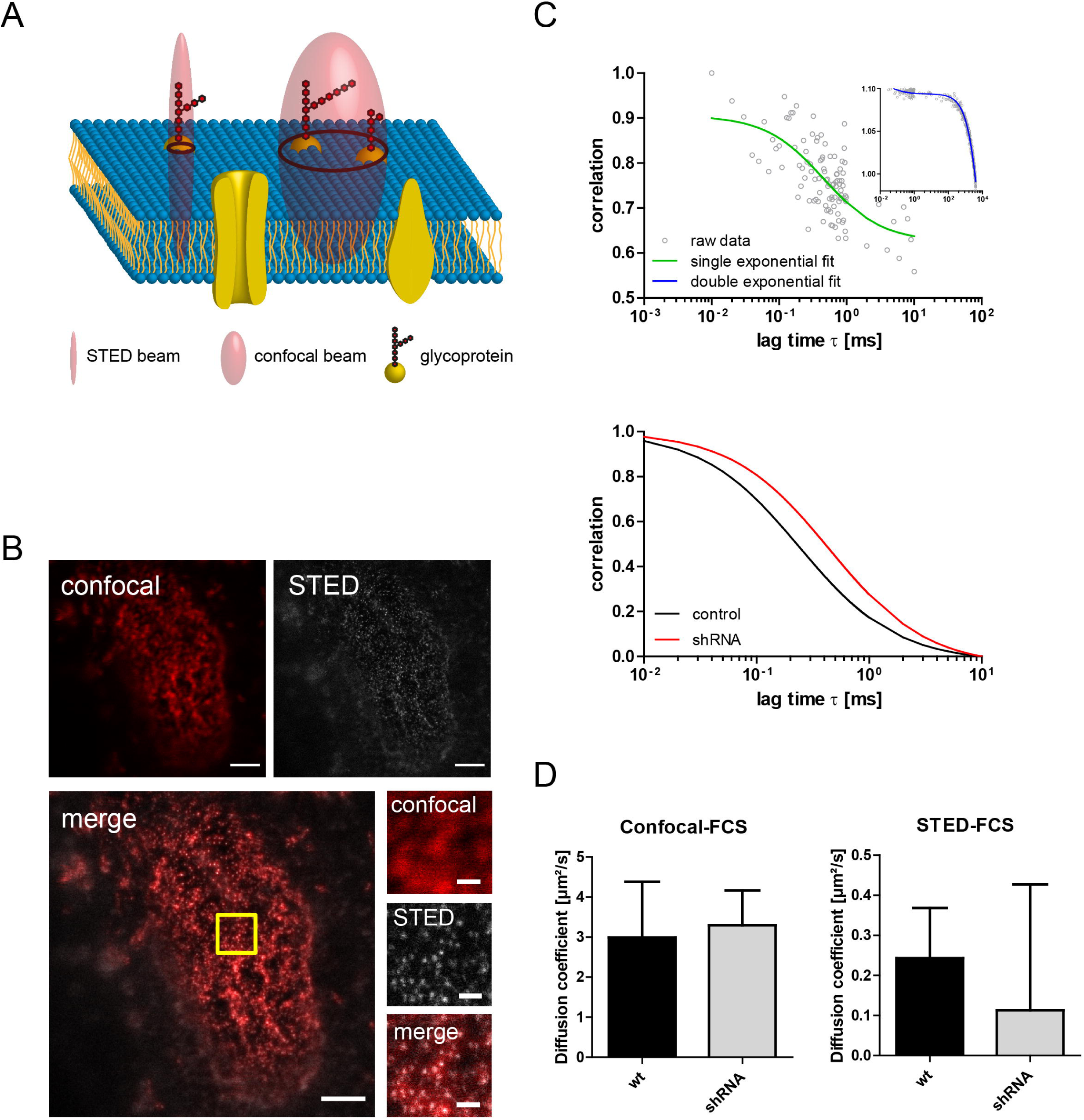
Characterization of carbohydrate chain-lengths of glycoproteins by STED-FCS. (A) Experiment schematic view: glycoproteins are located at the cellular plasma membrane, confocal-FCS excitation and detection are not selective, because many different carbohydrate chains are included in the same excitation spot. On the contrary, in the twenty time smaller FCS-STED excitation and detection volumes only few glycoproteins per location (down to an isolated one) can be excited and detected. (B) Top, overview of cell labelled with WGA-Atto647N and imaged in confocal and STED mode (left and right picture, respectively). Bottom left, overlay of the confocal and STED picture. Bottom right, detailed view from region of interest marked in yellow (from top to bottom confocal, STED and confocal plus STED merge). (C) Top, confocal-FCS and STED-FCS autocorrelation curves. (Inset) the full autocorrelation curve could be fit by a double exponential model function (blue line) as described in equation 4 (online Supplementary Methods). Single exponential model function (green line) was used to fit the first part of the correlation curve relative to lag times not exceeding 10 ms. Bottom, two typical autocorrelation curves fitted from a STED-FCS measurement on untreated cells (control, black lines) and shRNA treated cells (red line). A clear reduction of the diffusion coefficient in the case of shRNA treated cells can be already noticed. (D) Quantitative evaluation of the diffusion coefficients of shRNA-treated or wt cells by means of confocal-FCS (left) and STED-FCS (right). Confocal-FCS could not detect any difference in the diffusion behavior of wt and shRNA treated cells. STED-FCS detected a clear reduction of the average diffusion coefficient measured from shRNA-treated cells together with an enlargement of the distribution of the diffusion coefficients (see Table S1). Please note that diffusion coefficients are in general lower using the STED-FCS approach in comparison with confocal-FCS. This is due to anomalous diffusion as demonstrated by studying the diffusion coefficient tendency versus the variation of the excitation detection volume (for more details please see Supplementary Figure S2 online). Scale bars are 2.5 µm (overview scans) and 0.5 µm (regions of interest), respectively.

In Figure 2B (first row), the WGA-labelled cells were imaged in confocal (left) and STED mode (right). In Figure 2B, the overlay of confocal and STED images are shown. The square region of interest (ROI) depicted on the left corresponds to the magnified views shown on the right, comprising confocal microscopy, STED microscopy, and an overlay of both modalities. As shown in Figure 2B, STED imaging produced smaller clusters than confocal imaging. The average diameter of obtained clusters were 81 nm ± 21 nm. The diameter of the detected clusters, which defines the detection volume, is liable to modulation via the STED beam power. Elevated power levels could theoretically have facilitated the detection of volumes approaching that of an individual carbohydrate chain. However, to mitigate the potential phototoxic effects, the detection radius was maintained at approximately 80 nm.

In Figure 2C, the exemplary ACs of WGA-Atto647N diffusion are shown. This revealed that two diffusing species can be measured using a double exponential fit model (inset blue line) defined in equation 4 (online Supplementary Methods). FCS measurements were not set for accurate determination of the longer (>10s) diffusion times τ(d2) associated to almost static molecules. Our study was rather focused to determine the fast components of the diffusion curves. Therefore, only diffusion times τ(d1) <10 ms were evaluated by fitting a part of the autocorrelation function with the single component 3D diffusion model described by equation 2 (online Supplementary Methods). In the latest analysis, the maximal time interval τ, to which the signal is correlated (maximal lag time), was set to be 10 ms. In Figure 2C (bottom line) the normalized fit functions of two exemplary STED-FCS curves for shCTL-(black line) and shEXT1-treated cells (red line) are shown. This clearly indicates a shift towards shorter τ(d1) diffusion times upon treatment and resulting from shorter carbohydrate chains due to knockdown of EXT1. In Figure 3D, diffusion coefficients of the fast component [τ(d1)] were measured by confocal-FCS (on the left) and STED-FCS (on the right) in shCTL- and shEXT1-treated cells. The confocal-FCS measurements suggested that mean and dispersion of the diffusion coefficient remained unchanged in treated cells failing to differentiate treated and untreated samples. In contrast, STED-FCS analysis revealed clear differences between shCTL- and shEXT1-treated cells. These differences were observed in both the average diffusion coefficient and its dispersion. In both cell types, diffusion coefficients measured by STED-FCS were 10-fold lower, due to anomalous diffusion, than those measured by confocal-FCS. Next, comparing now only STED-FCS measurements on control and EXT1 knockdown cells, an average relevantly smaller diffusion coefficient is detected in the shEXT1 case. As a result, STED-FCS might be used as a tool to distinguish different stages of pathologies by detecting changes in both the average diffusion coefficient and its dispersion. These findings highlight a promising path for future applications of this STED-FCS methodology.

**Figure 3:**
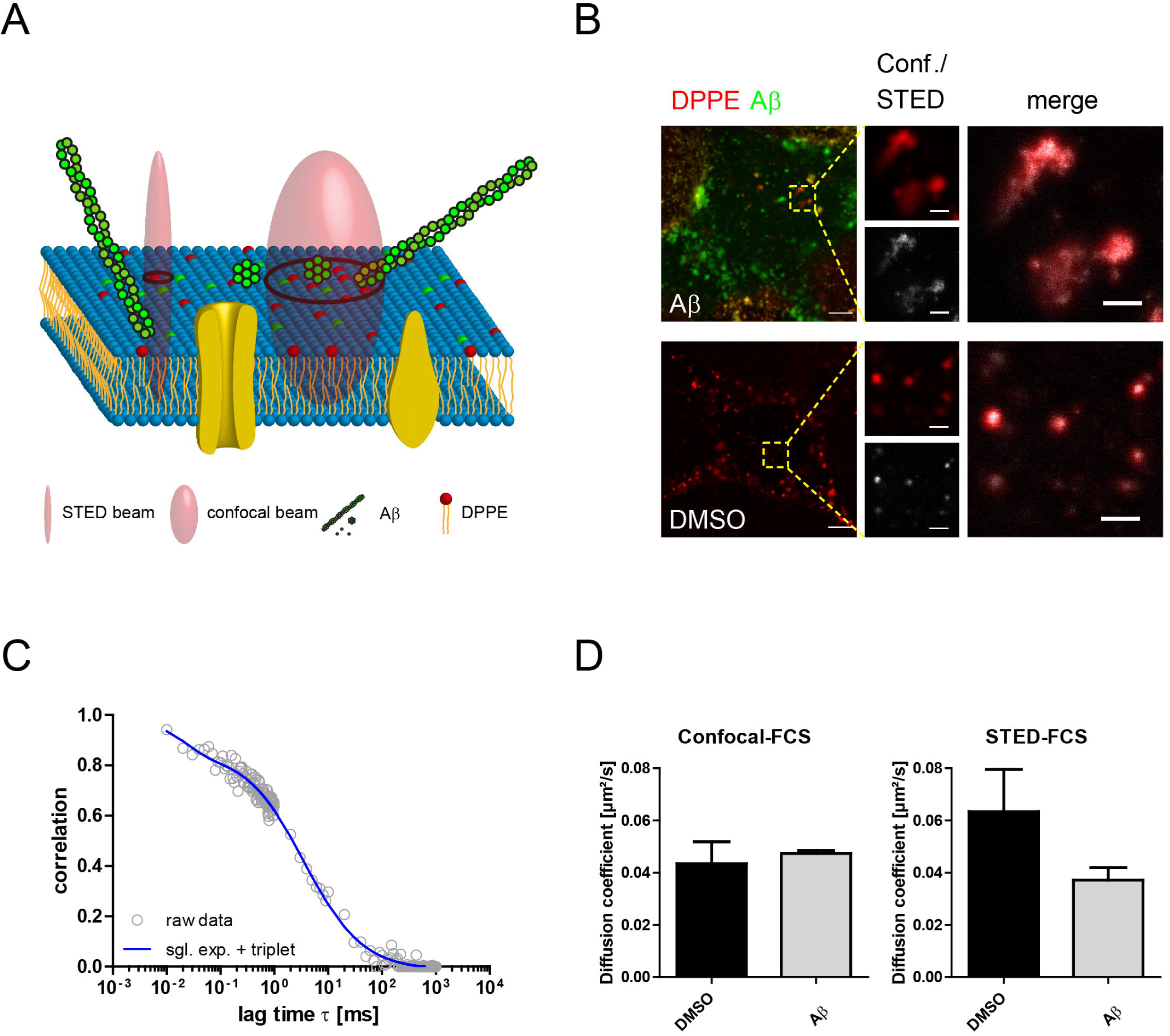
Alteration of membrane viscosity in Aβ42-treated cells. (A) Experiment schematic view: DPPE is a freely floating lipid analogue that was monitored by FCS when tagged to Abberior Star Red. Smaller clusters and fibrillary assemblies of the self-aggregating Alzheimer disease peptide Aβ42 alter the fluidity when bound to the membrane. The smaller STED-FCS excitation and detection volumes promise more confined measurements in the vicinity of Aβ clusters as opposed to confocal-FCS. (B) Top, confocal overview scan of DPPE-Abberior Star Red (red) and Aβ-FITC (Green, top) or DMSO vesicle control (bottom) in mHippo cells. Middle, DPPE signal from magnified region of interest in the overview scans marked in yellow, recorded in confocal (red) or STED (gray) mode. Right, merge of the two images from the middle panel. (C) Autocorrelation curves of DPPE could be fitted by a single component 3D diffusion model with triplet state contribution (see equation 5 in online Supplementary Methods). The triplet state is dye-specific and visible as a short decay at the beginning of the curve (<0.1 ms). (D) The diffusion coefficients were calculated for confocal-FCS (left) and STED-FCS (right). With confocal-FCS, no differences in the diffusion characteristics of DPPE were apparent after Aβ treatment (Aβ) compared to the vehicle control (DMSO). STED-FCS revealed a reduction of the diffusion coefficient by almost 50 % in Aβ-treated cells. DPPE is a normally freely diffusing lipid that should not be slowed down by interaction with endogenous molecules in the plasma membrane. Hence, either the direct interaction with Aβ in nanodomains or a locally decreased viscosity therein is slowing down the diffusion of DPPE. Scale bare are 5 µm (overview scans) and 1 µm (insets), respectively.

**Figure 4:**
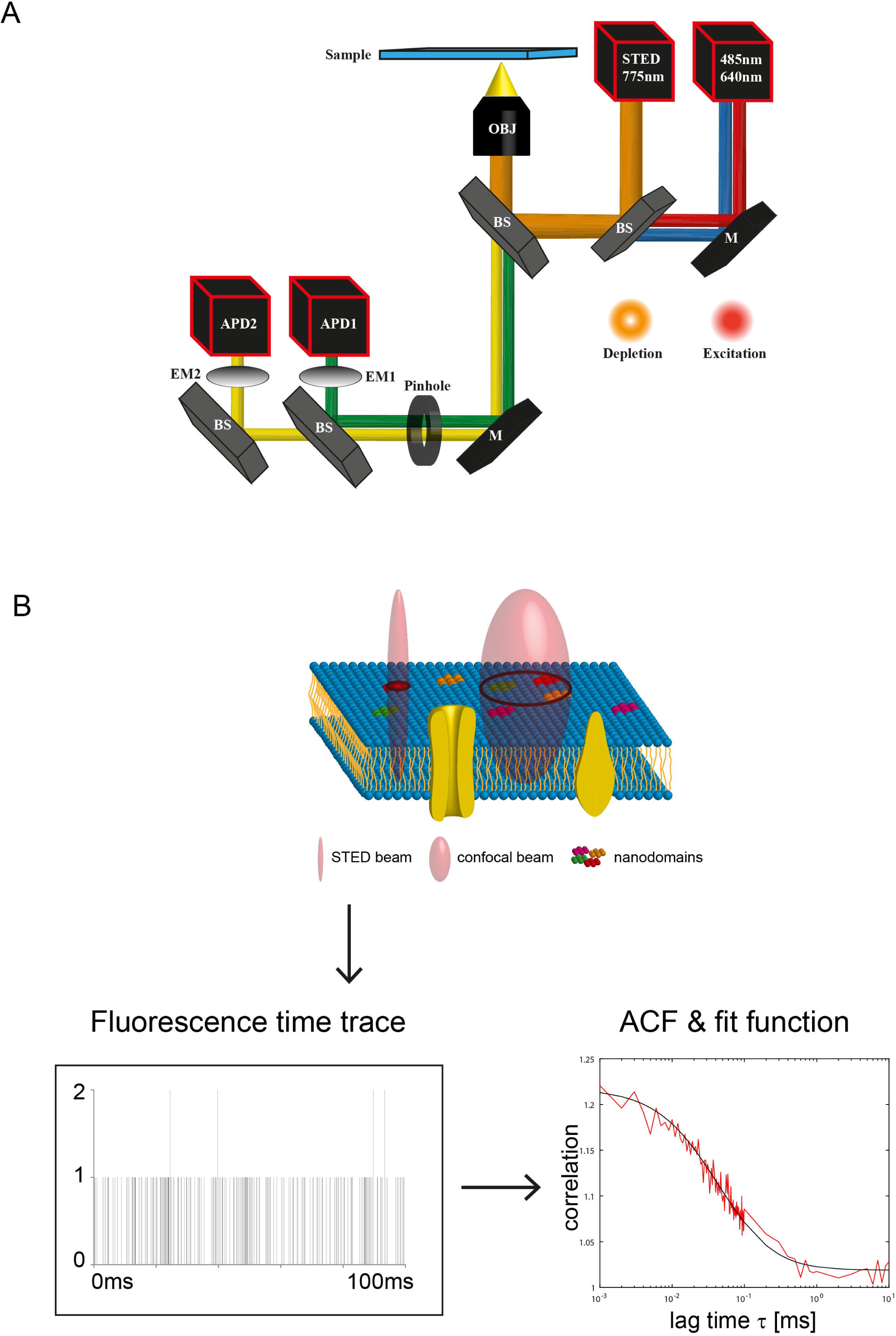
Setup of the STED microscope and outline of STED-FCS measurements in the plasma membrane of cells. (A) For STED-FCS measurements, an Abberior easy 3D STED “infinity line” system was used. For excitation two pulsed lasers emitting at 485 nm and 640 nm were employed. A pulsed beam emitting at 775 nm was used for depletion after a spatial light modulator formed the donut-shape. For the majority of experiments, except when stated differently, the 3 W depletion beam was run at 20 % power generating a transversal excitation radius of about 38 nm. The beams were directed via mirrors (M) and beam splitter (BS) through a Nikon Ti microscope body equipped with perfect focus system and a 60x P-Apo oil objective. The emitted light passed a pinhole set to 1 airy unit for all measurements, and was detected through GFP (495/25 nm, EM1) and Far-Red (685/70 nm, EM2) filters on independent avalanche photodiodes (APD 1&2). Abberior Imspector software was used to control the setup. BS = beam splitter. EM = emission filter. M = mirror. (B) The excitation beam is standing in a fixed position and fluorescence time traces were recorded every 1 µs (please note the that the relative size of the confocal/STED excitation beam is only qualitatively depicted). The autocorrelation function (ACF) was calculated in MATLAB and a fit function, depending on the chosen diffusion model, was applied to extract parameters such as diffusion time τ(d) and fraction size in multicomponent models.

### Alzheimer’s disease-associated Aβ42 peptide alters plasma membrane fluidity

Finally, we show how STED-FCS can have a significant impact on the research of Alzheimer’s disease. Causative and pathomechanistic processes as well as drivers of this multifactorial and progressive neurodegenerative disease to some degree are still a matter of debate [43]. However, a relevant factor and neuropathological hallmark is the increased production and marked extracellular deposition (‘plaques’) of a misfolding- and aggregation-prone peptide named amyloid-beta (Aβ), which is generated by consecutive proteolytical cleavages of a much larger amyloid precursor protein (APP). Aβ peptides exhibit variations in their structure due to different starting (N-terminal) and ending (C-terminal) points. Diffusible aggregates of the Aβ1-42 fragment, known for their rapid clustering, are considered especially toxic to synapses and neurons. This pronounced toxicity stems from their ability to detrimentally interact with receptors on nerve cell surfaces or directly with the cell membrane [44, 45]. Since interactions of Aβ assemblies with the plasma membrane and consequences of the latter on Aβ toxicity and aggregation are of critical interest [46], a STED-FCS imaging assay was used to investigate the short-term effects of pre-aggregated Aβ on lipid diffusion. mHippo cells, a mouse neuronal cell line, were treated for 2 hours with either vehicle control (DMSO) or FITC-labeled Aβ1-42 (Aβ-FITC). The assay employed a lipid analogue DPPE conjugated to fluorescent Abberior Star Red dye. The measurements were performed within 1 hour of labelling to reduce any significant effects of endocytosis.

In Figure 3A, the STED-FCS imaging assay is illustrated. Confocal and STED-FCS measurements were performed on the dorsal cell membrane to understand the role of Aβ aggregates on the diffusion of DPPE. The 25 times smaller excitation volume of STED-FCS acquisitions provided the ideal probing region for measuring diffusion coefficients of DPPE. In Figure 3B (top), STED/confocal-FCS images of Abberior Star Red-labeled DPPE (red) and FITC-labeled Aβ (green) are shown. The control cells (treated with DMSO as vehicle) are shown below (Figure 3B). It is notable that Aβ-FITC-treated cells appeared oligomeric and that fibrillary assemblies of Aβ were also DPPE-positive. The magnified images of the DPPE signal are displayed in the middle panel, using confocal (red) and STED (greyscale) acquisition modes. The right panel shows the overlay of both channels, highlighting the enhanced resolution and detail achieved with STED over confocal microscopy. Confocal and STED-FCS acquisitions were targeted at the stable clusters of DPPE. In Figure 3C, the AC of DPPE is presented. The latter was analyzed by fitting the raw data (circled curve in Figure 3C) with a model fit function (blue line) that aims to describe the 3D diffusion of a dye whose emission is also influenced by triplet state transitions (for more details see equation 5 in online Supplementary Methods). It is notable that this triplet state transition is dye-specific. The triplet state contribution is responsible for the fast decay observed at lag times shorter than 0.1 ms in DPPE conjugated to Abberior Star Red. The diffusion coefficients extracted by the fitted ACs are shown as bar graphs in Figure 3D, on the left the ones measured from confocal-FCS autocorrelation curves, and on the right the ones measured from STED-FCS. In contrast to STED-FCS, confocal-FCS was unable to detect differences in DPPE diffusion between Aβ-treated and untreated samples. In summary, STED-FCS, which revealed reduced membrane fluidity upon treatment of cells with Aβ, is an optically non-invasive method to measure the impact of such proteopathic conformers on cell membranes.

## Discussion

This work explores the potential of point-scanning STED-FCS in living cells through three proof-of-principle studies. We aim to demonstrate that STED-FCS is a valuable imaging technique for a wide range of cell biology experiments. By utilizing APD-derived photon traces, we achieve microsecond-scale detection, enabling STED-FCS measurements of triplet states and nanoscale volumes. This capability allows for the observation and interpretation of biological processes that were previously largely inaccessible. In this study, we did not fully exploit the high temporal resolution that certainly can be achieved by STED-but we focused on biologically relevant timescales, which typically range from milliseconds to seconds. The three examples presented here, spanning diverse areas of biomedical research, highlight the broad applicability of this technique. Future perspectives and technical considerations for these studies are discussed below.

### A shift in diffusion state observed for LC3b when entering synapses

As a first proof-of-principle, we investigated the diffusion characteristics of free cytosolic LC3b in axons and presynapses of rat hippocampal neurons. The application of STED-FCS revealed a reduced diffusion coefficient of the molecules inside presynaptic boutons contrary to the extra-synaptic/axonal regions. This effect, partly due to anomalous diffusion [34], indicates that LC3b has reduced mobility inside synaptic sites. Presynaptic terminals are small compartments in which organelles (i.e., synaptic vesicles), filaments, and machinery involved in neurotransmission are tightly packed, hence representing a crowded environment. The slower dynamics of LC3b at presynapses found in this study are therefore explained by the likelihood of higher molecular crowding found in these compartments. Importantly, the technical limitations of confocal-FCS as standing alone technique did not allow us to detect the reduction of the “fast” diffusion coefficient and the enhanced anomalous diffusion, hence blocking us to decipher these newer findings. The technical limitation was that the confocal detection spot (employed by confocal-FCS) was comparable to the size of synaptic region. On the contrary, with the STED-FCS detection spot being smaller than the size of synaptic region, this approach allowed us to measure a significant decrease in the number of fast diffusing molecules. Furthermore, STED-FCS measurements of LC3b dynamics revealed the existence of a “very-slow” diffusion species of LC3b in synaptic boutons. Considering the key role of LC3b conjugation to autophagosome membranes during autophagosome formation, it would be interesting to explore if this slow fraction relates to this process, and whether STED-FCS could be used to visualize LC3-positive autophagosome formation in small compartments in combination with other autophagy markers.

As a result of this study, STED-FCS can be applied to quantitatively measure the dynamics of LC3b molecules (and likely other proteins of interest as well) at the synaptic sites. Since, to our knowledge, this is the first time STED-FCS is used for comparing synaptic/non-synaptic sites, the overall results shown in Figure 1 describe a new method for the study of the molecular dynamics of autophagy proteins other neuronal compartments. Future experiments should address (i) how synaptic activity modulates the dynamics of LC3 and other autophagy proteins at boutons, (ii) its effect on the recruitment of these proteins into synaptic compartments, and (iii) how dynamic changes correlate with autophagosome biogenesis at these sites.

### Detecting biologically relevant alterations in carbohydrate chain length of glycoproteins

The second proof-of-principle experiment demonstrated the diagnostic potential of STED-FCS. The aim of this experiment is detecting the shortening of sugar chain length of glycoproteins. Again, confocal-FCS suffers from the large diffraction-limited detection volume, which results in the simultaneous detection of numerous carbohydrate chains. Consequently, this yields to the measurement of an averaged diffusion coefficient greatly reducing the sensitivity in capturing diffusion fluctuations. In contrast, the approximately 25 times smaller detection volume of STED-FCS enables the detection and measurement of diffusion coefficients for either individual or a fewer carbohydrate chains. Additionally, carbohydrate chains are located on the plasma membrane that has an approximate thickness of about 5-10 nm, thus, the effective detection volume is further reduced. Theoretically, the STED-FCS detection spot can be confined to even smaller volumes, however, consideration was given to photo toxicity. Therefore, we used a relatively low STED beam power to minimize any adverse effects on living samples. Hence, an optimal compromise between detection volume and low photo toxicity was achieved, while maintaining the feasibility of this method for living samples. STED-FCS measurements showed that the diffusion coefficient of the fast fraction of WGA-bound glycoproteins was higher in control cells compared with EXT1-knockdown cells. The smaller diffusion coefficient observed with EXT1-knockdown cells was attributed to anomalous diffusion of WGA-Atto647N. A similar observation was made during studying its diffusion coefficient tendency versus the variation of the excitation detection volume (Supplementary Figure S2). The wider range of diffusion coefficient values observed in cells treated with shEXT1 could be misinterpreted as showing no difference between shEXT1-treated and control cells. On the contrary, the large dispersion of the diffusion coefficient associated to the shRNA indicates that, during this treatment, a huge variety of different carbohydrate chains are formed by “cutting off” of the original one. In confocal-FCS measurements (on volumes that contain many carbohydrate chains) these dispersions are blurred out. Based on these findings, STED-FCS has demonstrated to be capable to measure the propagation of sugar-binding proteins and potentially the identification of pathologically altered glycoproteins in living biological samples. Recently, it has been suggested that the dynamic switching of heparan sulfate binding proteins between neighbouring sugar chains is central for the formation of protein gradients in tissues and, as such, crucial for tissue development and cell-cell communication [47]. In line with this, we measured a reduced diffusion of WGA on cell surfaces with shorter heparan sulfate chains. WGA was selected as a model protein; however, future studies applying STED-FCS will likely be suitable to measure the movement of other sugar-binding proteins on cells and in tissues.

### Aβ aggregates impact on membrane fluidity

The final comments relate to the third proof-of-principle experiment of STED-FCS applications. Murine hippocampal cells were subjected to Aβ, an aggregating and neurotoxic peptide increasingly produced in the aging brain and critically associated with development and progression of neurodegenerative and currently incurable Alzheimer’s disease. Our STED-FCS results have detected a slower diffusion of DPPE upon treatment with the aggregated Aβ peptide. The result implied that Aβ aggregates have not hindered the functionality of plasma membrane in our experiments. Given that DPPE, typically has autonomous diffusion and extreme low affinity for endogenous plasma membrane molecules. Confocal-FCS measurement however could not isolate them mainly because the size of the detection volume exceeds the size of the Aβ clusters. Thus, the measurements of changes in plasma membrane viscosity are challenging to be performed with confocal-FCS as previously demonstrated by Justin and colleagues [48]. However, STED-FCS measurements (by reducing circa 25 times the excitation/detection volume) measured a two times smaller diffusion coefficient of DPPE in presence of Aβ. These results indicate that the presence of Aβ, has reduced the diffusion speed of DPPE. From our data, it is still not fully clear if Aβ slowed down DPPE by directly interacting with it or by locally reducing membrane viscosity. Our experiments let us guess that Aβ locally reduced membrane viscosity, because this was the only experiment presented in this paper were we did not detect anomalous diffusion, which in general is generated by strong molecular interactions. A clear answer of this question, however, will need further and more detailed investigations. Whatever the actual reason is, STED-FCS measurements performed on DPPE showed uncontrovertibly that cell membrane functionality is affected by Aβ aggregates.

## Conclusions

This study employed STED-FCS as a powerful tool to observe protein dynamics in living cells, demonstrating its broad applicability across diverse biological studies. STED-FCS enabled visualization of molecular diffusion within nanoscopic cellular compartment generating sub diffraction limit excitation volumes. We examined protein interactions in synapses, conformational changes in glycoprotein modifications, and a potential alteration in neuronal membrane fluidity upon treatment with Alzheimer’s disease-associated Aβ. These findings may inspire the development of alternative methods for validating biochemical assays and even contribute to advancements in sub-diffraction-limited correlation spectroscopy. Given the increasing presence of STED microscopy systems in research laboratories and imaging core facilities, we aim to lower the barrier to STED-FCS adoption by showcasing its practical application to a wide range of biological research questions. Our goal is to encourage its wider use in investigating the detailed mechanisms of protein interactions and membrane-associated processes.

## Supporting information

Supplementary Material_FCS

## Acknowledgements

We thank Rukan Nasri from the Institute for Nanostructure and Solid State Physics of the University Hamburg for providing the chamber slides for STED-FCS diffusion calibration. We also thank Neeraj Prabhakar, now self-employed scientist, for reviewing the manuscript.

