## Supplementary Material_FCS for "STED-FCS in subdiffraction limit volumes reveals altered diffusion in live cell applications"

### Supplementary Methods

#### Data analysis

Using Matlab software, fluorescence time traces were imported, binned to 10 µs steps and at first the autocorrelation function (ACF) was computed according to the following equation:

$$G(\tau) = \frac{\langle F(t) * F(t+\tau) \rangle}{\langle F(t) \rangle^2} \qquad \text{\{equation. 1\}}$$

where F is the fluorescence signal, t is the time point when the signal was recorded, and τ is the so-called lag time, a delayed time point in the time trace to which the signal is correlated in the calculation of the ACF. A least squares fit was performed on the ACF using different adaptation functions that are described by the equations below. For reducing computational load, not all data points (N) of the ACF were used but for N<100 every single data point, for N=100-1000 every 100th data point, and for N>1000 every 1000th data point.

Depending on the experiment, different adaptation function each representing a certain diffusion models were used to fit the ACF:

For the experiments described in Figure 2 and Figure 3, a 3D diffusion model was chosen to fit the first 10-20 ms of the ACF in order to retrieve the fast component τ(d1):

$$G = Off + Amp * \left( 1 + \frac{\frac{1}{N(P)} * 1}{\left(1 + \frac{t}{\tau(d1)}\right)^{\alpha}} \right) * \left(\frac{1}{\sqrt{1 + \frac{t}{\tau(d1) * k^2}}}\right) \qquad \text{\{equation. 2\}}$$

Off is an offset and was set to 1. Amp is the amplitude of the fit function, and N(P) is the number of diffusing particles. Please note that, $\kappa = \kappa_{conf}$ and $\kappa = \kappa_{STED}$ are the beam waist ratios of the confocal and STED beams, i.e., the quotient of the axial and lateral radii of the detection volume measured with sub-resolution fluorescent beads. $\kappa_{conf}$ and $\kappa_{STED}$ were determined to be 2.2 and 12 (for 20 % STED beam power), respectively. α is the anomalous factor that describes the degree of hindered diffusion and was kept in the range of 0.8-1 according to [1]. The transit time of the fast diffusing species [τ(d1)] was extracted and the diffusion coefficient D calculated according the formula:

$$D = \frac{r^2}{4*\tau(d)} \qquad \text{\{equation 3\}}$$

where $r_{conf}$ and $r_{STED}$ are the radii of the confocal and STED detection volumes and were determined to be 133 nm and 38 nm, respectively.

The ACF of the full-length time traces in the LC3b experiments (Figure 2) also revealed an additional, slower diffusing species in the seconds regime. For fitting the curve, the following two components 3D diffusion model was employed:

$$G = off + Amp * \{1 + \frac{1}{N(P)} * \left[ \left( \frac{1}{\left(1+\frac{t}{\tau(d1)}\right)^{\alpha}} \right) * \left( \frac{F1}{\sqrt{1+\frac{t}{\tau(d1)*k^2}}} \right) + \left( \frac{1}{\left(1+\frac{t}{\tau(d2)}\right)^{\alpha}} \right) * \left( \frac{1-F1}{\sqrt{1+\frac{t}{\tau(d1)*k^2}}} \right) \right] \}$$

{equation 4}

τ(d1) was previously determined with the single component model (equation 2) and a shortened ACF (lag times up to 100 ms). This value was then kept as a constant for fitting the full length ACF (lag times up to 60 s) of the same fluorescent time trace with equation 4. This allowed to extract the fraction of fast diffusing molecules (F1) and theoretically the transit time of the slower component [τ(d2)]. However, an accurate determination of τ(d2) would have required minutes of recording time [2], so only F1 was extracted. The values are summarized in Table S1.

For the experiment described in Figure 3, a single component 3D diffusion model with triplet state extension was used:

$$G = Off + Amp * \left[ 1 + \frac{1}{N(P)} * \frac{1}{\left(1+\frac{t}{\tau(d1)}\right)^{\alpha}} * \left( \frac{1}{\sqrt{1+\frac{t}{\tau(d1)*\kappa^2}}} \right) * \left( 1 + \frac{T}{1-T} * e^{-\frac{t}{\tau(T)}} \right) \right] \qquad \text{\{equation 5\}}$$

This was necessary to fit the first slope of the ACF in the 0.1 ms regime, which resembles a dye-specific triplet state conversion. Here, T is the fraction of molecules in the triplet state and τ(T) the triplet state correlation time. The diffusion time τ(d1) was extracted and the diffusion coefficient calculated as before.

#### Figures and Tables Legends

**Figure S1:** Spot size calibration.

The effective spot size of the 640 nm excitation beam was determined setting the 775 nm depletion beam to for five different intensities, ranging between 0% (confocal-FCS) and 20% (STED-FCS). Single far-red fluorescent 36nm diameter beads were used for the measurement series. Full width at half maxima (FWHM) of the excitation spots were determined and plotted against the STED depletion beam power.

**Figure S2:** Diffusion characteristic of WGA-Atto647N in MV3 cells.

MV3 cells were treated with non-targeting (wt) or shRNA (shRNA) that effectively shortens the carbohydrate chains in glycoproteins. Cells were labelled with WGA-Atto647N and the diffusion coefficients measured using STED-FCS for different STED beam powers in the range of 0-20 %. The diffusion coefficients were plotted against the equivalent spot diameters of the detection volume revealing anomalous diffusion of WGA-Atto647N in the samples. A decrease of the diffusion coefficient with smaller spot sizes in both conditions is the evidence for general anomalous diffusion of WGA-ATto647N in MV3 cells.

**Figure S3:** Diffusion coefficient vs. spot size in a freely diffusing sample.

The diffusion coefficients of Atto647N-labelled IgG antibodies were measured for different spot sizes in solution. The 775 nm STED depletion beam power was tuned in a range between 0.5 % and 20 % and plotted against diffusion coefficient. The measured values were constant in the observation range between, as expected for a freely diffusing species. Differences observed in the outlined live cell experiments have biological reasons and were not affected by the STED laser beam itself. Values are mean ± standard deviation for the four measurements.

**Table S1:** Results of the confocal and STED-FCS measurements in live cells as outlined in Figure 1- Figure 3. All values are harmonic mean ± SEM except for fractions which are mean ± SEM.

| | | Diff. coefficient [μm²/s] | Fraction (fast component) |
|---|---|---|---|
| **LC3-SNAP-SiR in hippocampal neurons** | | | |
| **STED-FCS** | LC3b - Synaptic site | 0.41 ± 0.16 | 18 % ± 3 % |
| | LC3b - Non-synaptic site | 0.70 ± 0.14 | 26 % ± 3 % |
| | Control - Synaptic site | 2.93 ± 1.04 | 10 % ± 2 % |
| | Control - Non-synaptic site | 3.31 ± 0.96 | 16 % ± 4 % |
| **Confocal-FCS** | LC3b - Synaptic site | 1.94 ± 0.15 | n.d. |
| | LC3b - Non-synaptic site | 1.97 ± 0.3 | |
| | Control - Synaptic site | 7.69 ± 0.90 | |
| | Control - Non-synaptic site | 6.52 ± 0.94 | |
| **WGA-Atto647N in MV3 cells** | | | |
| **STED-FCS** | wt | 0.24 ± 0.12 | n.d. |
| | shRNA | 0.11 ± 0.31 | |
| **Confocal-FCS** | wt | 2.99 ± 1.39 | |
| | shRNA | 3.29 ± 0.87 | |
| **DPPE-Abberior Star Red in mHippo cells** | | | |
| **STED-FCS** | + DMSO (control) | 0.063 ± 0.016 | n.d. |
| | + Aβ 1-42 (in DMSO) | 0.037 ± 0.05 | |
| **Confocal-FCS** | + DMSO (control) | 0.043 ± 0.008 | |
| | + Aβ 1-42 (in DMSO) | 0.047 ± 0.001 | |
