## Supplementary figures and images for "STED-FCS in subdiffraction limit volumes reveals altered diffusion in live cell applications"

### Figure S1

#### Slide 1
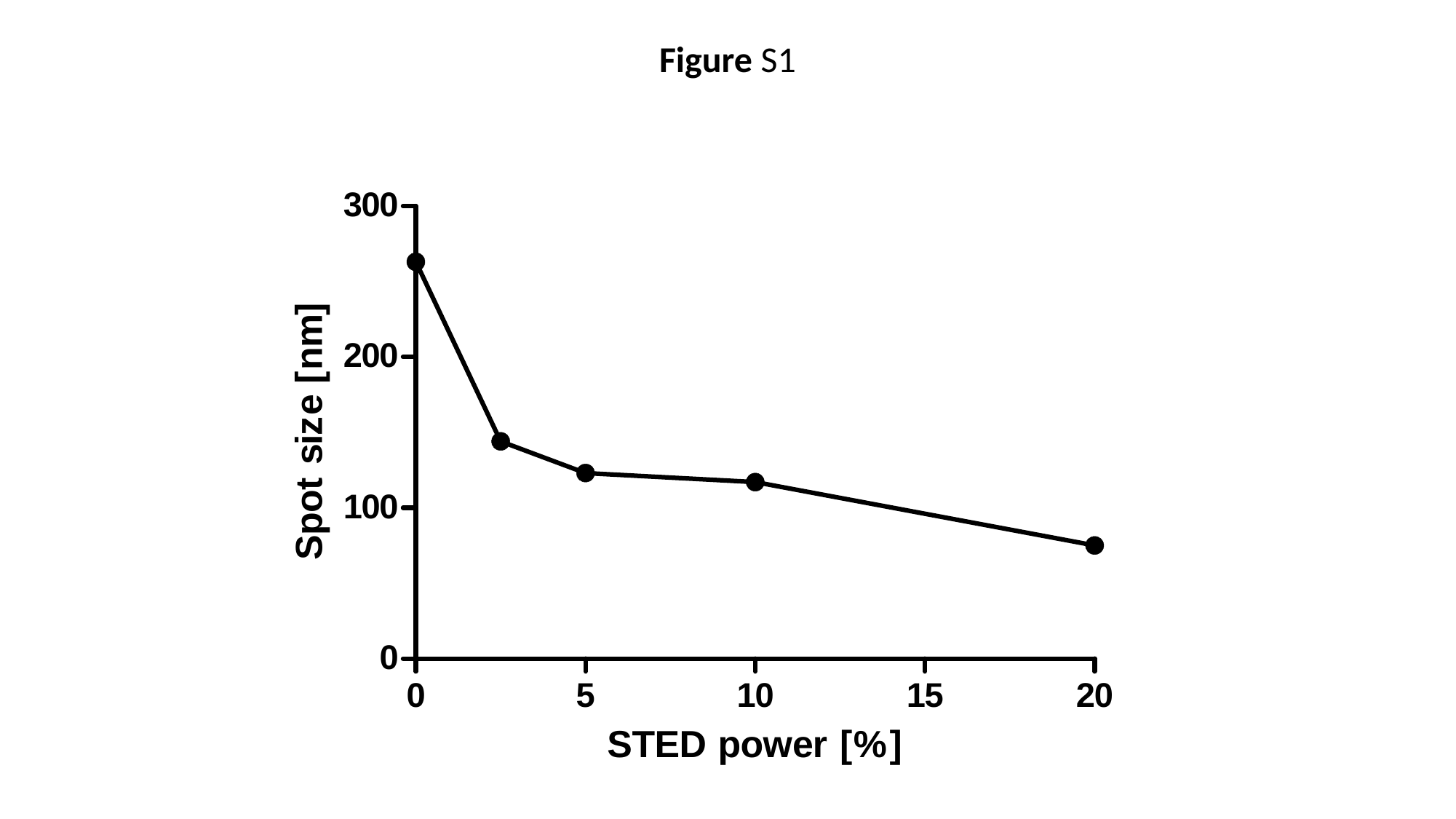

Figure S1

### Figure S1

#### Slide 1
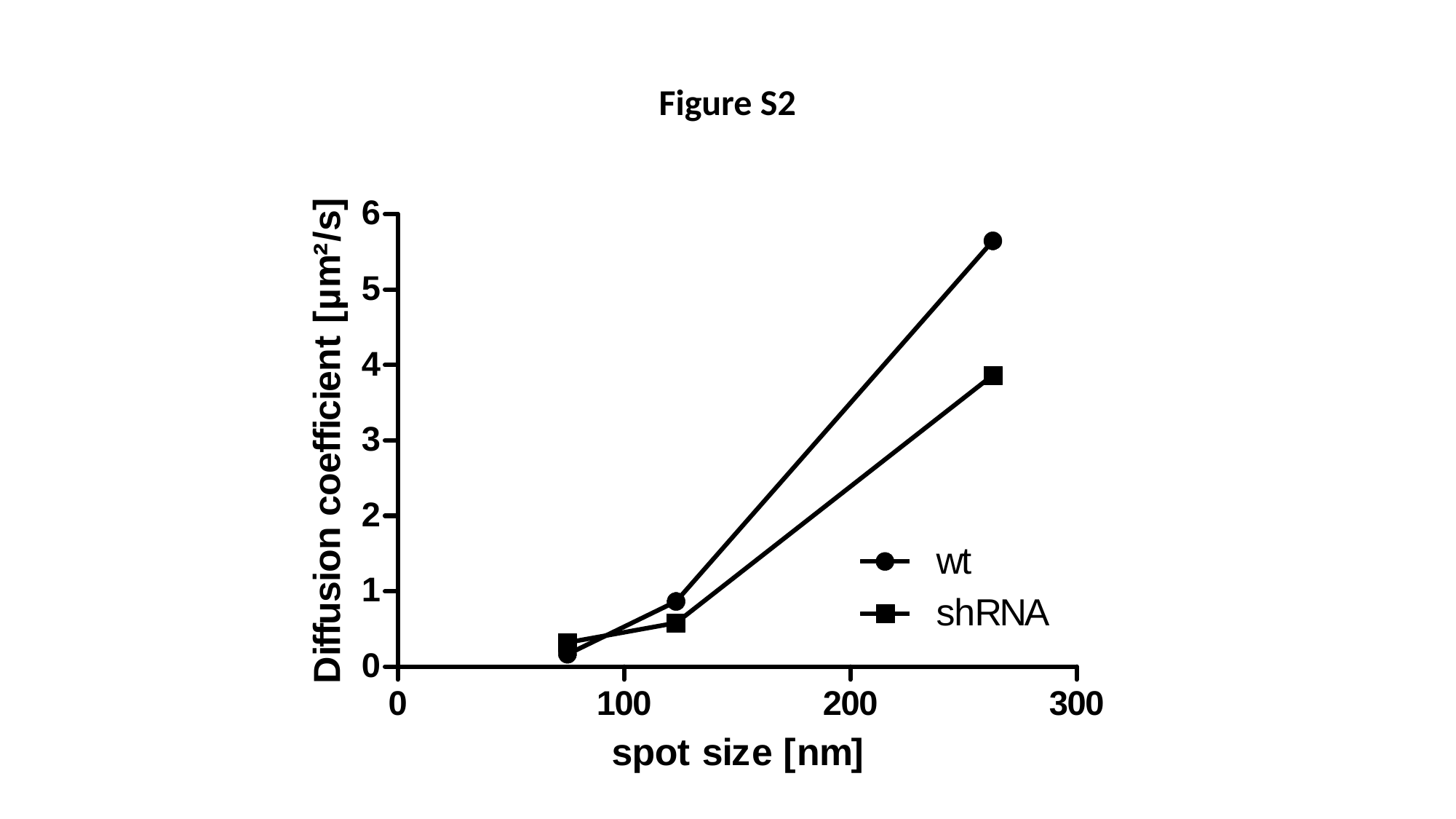

Figure S2

### Figure S1

#### Slide 1
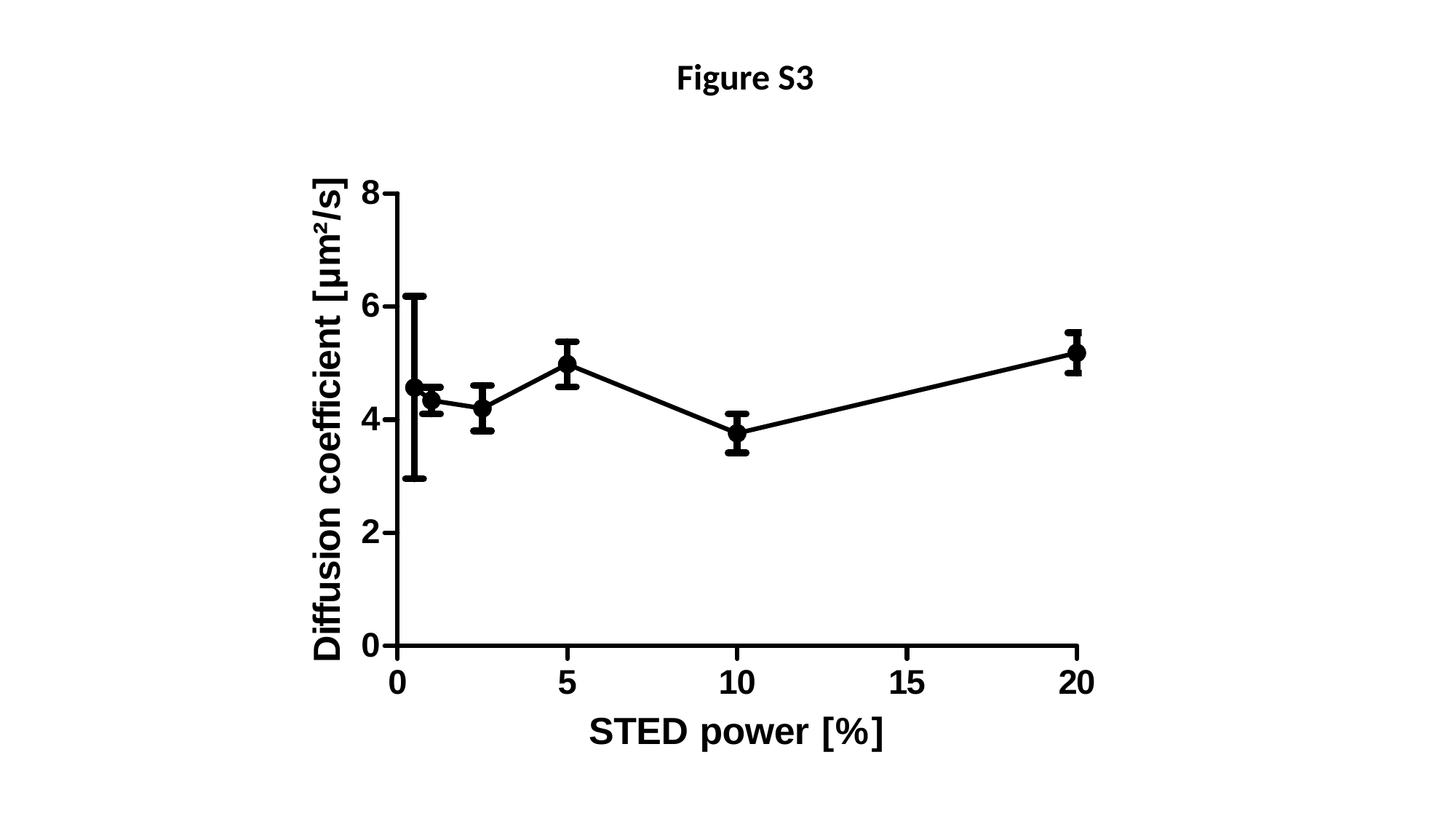

Figure S3
